# Machine Learning-Driven Phenotype Predictions based on Genome Annotations

**DOI:** 10.1101/2023.08.11.552879

**Authors:** Janaka N. Edirisinghe, Samaksh Goyal, Alexander Brace, Zahmeeth S. Sakkaff, V. S. S. L. Karanam, Dileep Kishore, Ricardo Colasanti, Tianhao Gu, Boris Sadkhin, Qizhi Zhang, Roy Kamimura, Christopher S. Henry

## Abstract

The rapid expansion of isolate genome sequences and metagenome-assembled genomes has created a growing need for computational approaches that can infer microbial phenotypes directly from genome information. Here, we present a machine learning framework that uses RAST-derived functional annotations as predictive features to classify bacterial phenotypes. We evaluated six machine learning algorithms; K-Nearest Neighbors, Gaussian Naive Bayes, Support Vector Machines, Neural Networks, Logistic Regression, and Decision Trees across three phenotype-prediction tasks: Gram-stain type (Gram-negative and Gram-positive), respiration type (aerobic, anaerobic, and facultative anaerobic), and growth phenotype (growth and no growth). All six classifiers showed strong predictive performance, although performance varied by phenotype. Gram-stain type was predicted with near-perfect performance, with precision and recall exceeding 0.99 across classifiers. Respiration type was predicted with a maximum overall accuracy of 81.5% using Gaussian Naive Bayes, while growth versus no growth on individual carbon substrates reached approximately 86% accuracy. Decision-tree models additionally provided interpretable links between phenotype predictions and specific functional roles, enabling biological interpretation of the features underlying classification. To make the approach broadly accessible and reproducible, we implemented a four-application workflow in the U.S. Department of Energy Systems Biology Knowledgebase (KBase) that enables users to: (i) upload high-quality data to train classifiers; (ii) annotate genomes in the training set using the RAST annotation algorithm; (iii) build six different genome classifiers; and (iv) predict the phenotypes of unclassified genomes. Together, these results demonstrate that genome annotations provide an effective and interpretable feature space for machine learning-based microbial phenotype prediction and establish a reusable framework for extending this approach to additional microbial traits.

## Introduction

Over the last two decades, we have witnessed an exponential growth in the number of genome sequences available, including over 300K isolate genomes in PATRIC (Wattam et al. 2017) and now millions of metagenome-assembled genomes (MAGs) spread across numerous collections (Nayfach et al. 2021). The recent rapid growth in MAGs in particular exposes a new trend in genomics where most of the genomes we are analyzing today have never been isolated or cultured in a lab (Parks et al. 2017). Experimental approaches for studying the physiology of microbial species have failed to keep pace with the emergence of new genome sequences, and such approaches are also impossible to apply to species that have not been isolated or cultured in the lab. Yet, many analyses that could be applied to understand and engineer the behavior of individual species, microbial communities, and even complex ecosystems require some knowledge of species physiology that often cannot be acquired experimentally. For example, the accurate reconstruction of a metabolic model for a new species, and thus the accurate prediction of metabolic engineering applications and modifications, requires knowledge of the growth conditions, gram stain, and respiration type (Thiele and Palsson 2010).

With all of these developments in the field, it has become increasingly vital to develop computational approaches that leverage extensive compendia of experimental data collected for known reference organisms to predict the physiology of new species based on sequence data alone. Many recently published studies demonstrate the wide-ranging potential of machine learning (ML) to address this emerging challenge (Irsoy et al. 2012). This includes the use of ML to predict potential substrates for promiscuous enzymes (Pertusi et al. 2017), to identify genes responsible for antimicrobial resistance based on observed DNA mutations (Davis et al. 2016), and to predict cell phenotypes (Feldbauer et al. 2015; Jensen and Ussery 2013).

Here we demonstrate a new approach for applying machine learning to predict bacterial cell phenotypes from genome sequence, as shown in Figure 1. To demonstrate this approach we are using Gram stain, bacterial respiration types (aerobic, anaerobic and facultative) and phenotype growth types as example phenotypes for prediction. We selected these specific phenotypes as case studies because: (i) these are key phenotypes to support automated construction of high-quality metabolic models and thus predictions derived from this approach will be a key component in the ModelSEED automated model reconstruction pipeline (Henry et al. 2010; Faria et al. 2023); and (ii) ML has been previously applied to predict these phenotypes, providing an excellent basis for comparison and evaluation of our approach (Feldbauer et al. 2015; Jensen and Ussery 2013).

**Figure 1.**
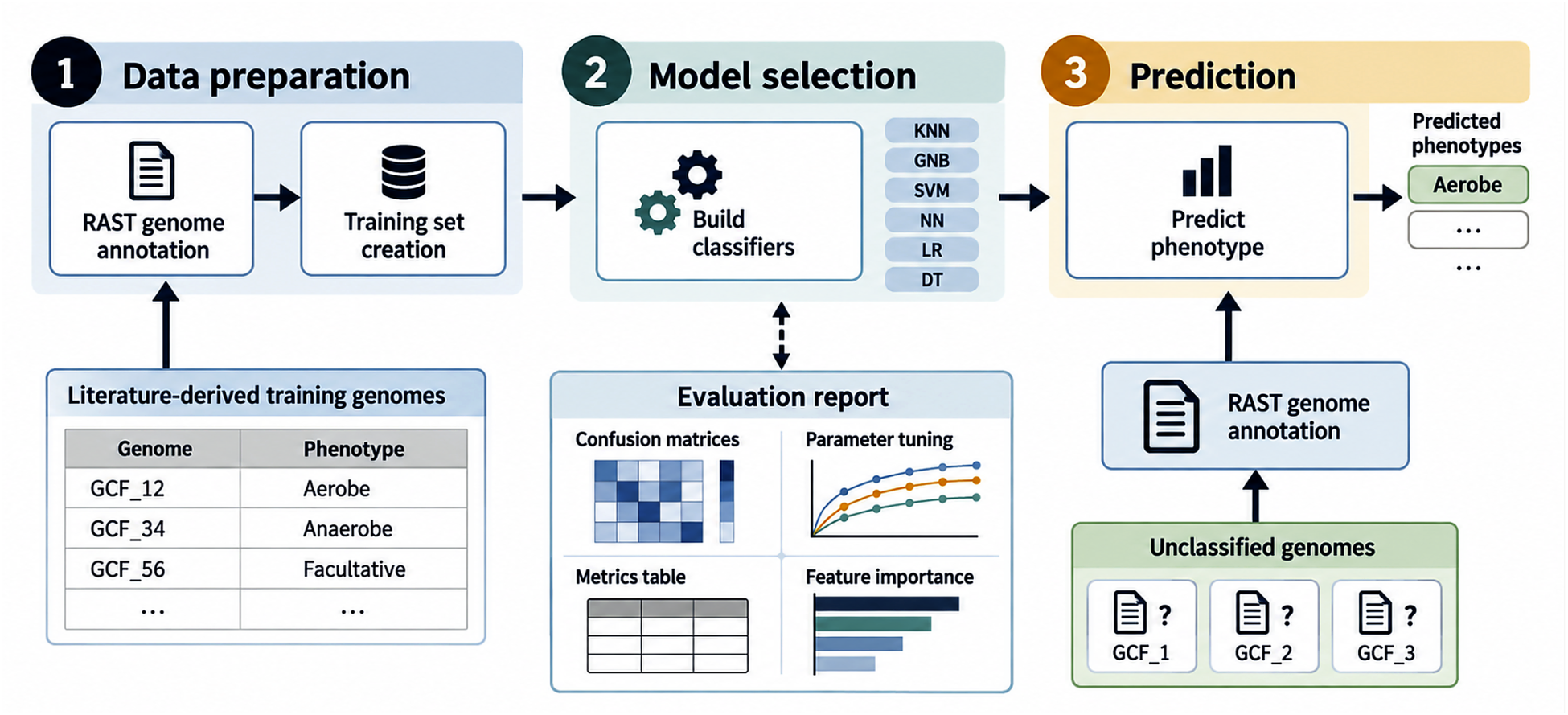
Machine learning based phenotype prediction. Literature derived genomes are annotated with the RAST annotation algorithm, and the respective phenotypes are used to build training sets. Based on the training sets, six classifiers are produced, namely, K-Nearest Neighbors (KNN), Gaussian Naive Bayes (GNB), Support Vector Machine (SVM), Neural Network (NN), Logistic Regression (LR), and Decision Tree (DT), to predict phenotypes (e.g., Gram or respiration type) for unclassified genomes.

One of the primary ways in which our new approach differs from previous ML efforts is in the choice of predictor used by our classifier to call phenotypes. In previous similar studies, phenotypes were called using protein families as predictors, e.g., Pfam and Clusters of Orthologous Groups (COG) (Feldbauer et al. 2015; Jensen and Ussery 2013). While such predictors are clearly effective for calling phenotypes, protein families have substantial drawbacks in this application. For instance, genes coding for promiscuous enzymes are able to catalyze multiple enzymatic transformations. In addition, families can lack precision as the families diverge, with the same family often mapping to many genes with significantly different functions. Conversely, many genes belonging to different families may perform the same function (de Crécy-Lagard et al. 2022). This is problematic, because the fundamental entity that defines a phenotype is gene function and this entity does not map cleanly to protein families. To avoid these problems, we selected the functions themselves as the predictors in our classifiers. The major challenge in this approach is that gene annotations are typically simple human-readable terms that describe the function of the gene, e.g., *ATP synthase alpha chain (EC 3.6.3.14)*. However, these strings do not conform to a strict, controlled vocabulary, making it difficult to apply ML to this problem. Thus, we specifically use SEED functional roles in our classifiers, as SEED manages this controlled vocabulary (R. Overbeek 2005) and provides a service, RAST (Brettin et al. 2015; Ross Overbeek et al. 2014), for rapidly assigning functions to new genomes based on sequence data. At the same time, we also acknowledge that other annotation ontologies like enzyme classes (EC) or gene ontology (GO) terms would likely work equally well as RAST functions. Another distinctive feature of this study is our exploration of a wider set of ML models (K-Nearest Neighbors, Gaussian Naive Bayes, Support Vector Machines, Neural Networks, Logistic Regression, and Decision Trees), whereas previous works focused exclusively on Bayesian networks (Jensen and Ussery 2013) and Support Vector Machines (Feldbauer et al. 2015). In contrast, classification techniques like decision trees are open to interrogation because they are composed of simple boolean rules that link predicted phenotypes directly to gene function. Here we will examine the rules proposed by our decision tree classifiers to explore the biological insights they reveal. For our two selected case-studies (respiration and Gram stain), we have assembled a phylogenetically diverse set of genomes as training data consisting of 2106, 549 for Gram stain and respiration respectively. We applied RAST (Ross Overbeek et al. 2014) to obtain consistent genome annotations for each of these genomes, and based on the literature, we have labeled each species with the known Gram stain type (Hucker and Conn 1923) and respiration type (Mazurie et al. 2010). We trained our six selected classification algorithms on this data to predict the target phenotypes.

## Methods

### Data Set Selection

The genomes that are used as the data set in this study were carefully derived from a phylogenetically diverse set of genomes. Respiration types were classified as aerobic, anaerobic, and facultative based on literature derived knowledge. Similarly, Gram stains were classified as Gram-negative and Gram-positive. Phenotypic growth data were classified as grow/no-grow for corresponding genomes and carbon sources. All the genomes were annotated using the RAST annotation pipeline and the functional annotations (functional roles) produced for each genome were used to build an attribute matrix denoting the presence or absence of each role. These attribute matrices served as the input feature space for our classifiers.

The three case studies discussed in this manuscript have considered the following datasets.

*Respiration type prediction* - The dataset for the respiration prediction task contained 549 genomes, of which 253, 126, 170 were labeled as aerobic, anaerobic, and facultative, respectively. The attribute matrix containing the presence or absence of each functional role had 549 rows (genomes) for the respiration data set and 44498 columns (functional roles).

*Gram stain prediction* - For this task we have collected 2106 genomes, where 847 and 1259 genomes are labeled as Gram-negative and Gram-positive, respectively. The attribute matrix for the Gram staining data set had 2106 rows and 36205 columns.

*Phenotype growth prediction* - The data set for phenotype growth prediction task contained 178 genomes and 64 substrates, where different genomes are labeled as growth or no growth for each of the substrates. Similarly, the attribute matrix for the phenotype growth data set had 178 rows (genomes) and approximately 30282 columns (functional roles). See Appendix A2 for links to the KBase Narratives containing the above explained data.

### Machine Learning Algorithms and Packages

The Python based scikit-learn (Pedregosa et al. 2011) package (version 0.23.2) was used to implement all the classification algorithms in this study. The output prediction targets are Gram stain and respiration type where the input features are functional roles derived from RAST annotation. Given the clear presence of class labels in our prediction tasks, our approach relies on supervised ML algorithms. In this study, we compare and contrast the outcome of six classification algorithms, namely, K-Nearest Neighbors (KNN), Gaussian Naive Bayes, Support Vector Machines, Neural Networks, Logistic Regression, and Decision Trees. For a more detailed description of the ML algorithms used in this work, refer to (Bishop 2006).

### Classification Algorithms

Out of the six popular classifier algorithms used in this study, KNN (Fix and Hodges 1989) was chosen to evaluate whether the high-dimensional attribute matrix derived from the annotation process intrinsically clusters the genomes by their biological properties such as respiration and Gram type, allowing for direct and accurate predictions. Gaussian Naive Bayes was chosen as it relies on conditional probabilities and could be used to find associative links between the many bacterial functional roles. Support Vector Machine with a radial basis kernel function was selected due to its effectiveness in high-dimensional learning settings where the number of features (here functional roles) is larger than the number of examples (Chang and Lin 2011). We chose to evaluate a standard Multilayer Perceptron Neural Network for its ability to learn complex nonlinear decision boundaries using several hidden layers composed via nonlinear activation functions. Logistic Regression was selected as it is often used in binary classification problems such as our Gram stain data and can be applied to multiclass classification settings like our respiration data through a one-vs-rest heuristic strategy. Lastly, the decision tree was selected because the functional roles being used to construct the decision nodes can be extracted and directly interpreted to glean scientific understanding by identifying the most relevant features in the data for the given prediction task. All classifiers were constructed with default parameters (Appendix A1) given by the scikit-learn implementation of the algorithms. The decision tree and the logistic regression algorithms contain random number generators; these were set with the same random seed (0) for all experiments.

### Decision Tree Tuning

As mentioned previously, the decision tree is a helpful algorithm since its internal mechanisms can be closely examined to interpret its classification path. We further optimized the decision tree classifier algorithm based on two parameters: tree depth and split criterion (Gini impurity, or Entropy) in order to reduce the risk of overfitting. After optimization, the decision tree with the highest test accuracy was analyzed to elucidate the most important functional roles implicated in solving a given prediction task.

### Training and Testing Sets

To evaluate our models, we followed a cross-validation procedure with ten training and testing folds using the StratifiedKFold scikit-learn module. Stratification ensures that the proportion of genomes with a specific classification in a fold remains similar to the proportion of genomes within the whole dataset. It is important to note that the ten training and testing folds were created prior to the construction and testing of the classifiers. Thus, all classifiers were presented with identical training and testing folds. The stratified fold method utilizes a random number generator which was set with the same random seed (0) for all experiments.

### Evaluation of Predictive Performance

To visualize the performance of each algorithm, we generated a confusion matrix (Ting 2017). The rows reveal the actual classification and the columns show the predicted classification. The main diagonal indicates the accuracy of each classifier, as it counts how many samples were predicted that actually had the same ground truth classification. The off diagonal elements indicate the false positives and false negatives. The percentage in each cell of the confusion matrix represents the average of that prediction over the ten cross-validation folds. To compare each of the classifiers we use the confusion matrix derived performance measures: accuracy, precision, recall, and F1 Score. Accuracy is the proportion of samples that are correctly classified. Precision is the proportion of samples predicted to belong to a particular class that truly belong to that class, whereas recall is the proportion of samples in that class that are correctly identified by the classifier. The F1 score is the harmonic mean of precision and recall and provides a balanced measure of classifier performance, with values closer to 1.0 indicating better overall performance.

### KBase applications and Source code

The source code related to the tools developed as part of this paper is provided in Appendix A2. The KBase application pipeline is comprised of four applications (i) Upload high-quality data to train classifiers (Supp. Figure 1); (ii) Annotate genomes in the training set with the RAST annotation algorithm (Supp. Figure 2); (iii) Build six different genome classifiers (Supp. Figure 3); and (iv) Predict the phenotype of unclassified genomes (Supp. Figure 4). The source code for the implementation of these applications is available in the GitHub repository: https://narrative.kbase.us/#catalog/modules/kb_genomeclassification. The three cases studies of Gram-stain (https://narrative.kbase.us/narrative/153575), prokaryotic respiration (https://narrative.kbase.us/narrative/153678) and phenotype growth prediction (https://narrative.kbase.us/narrative/ws.137002) were conducted in the KBase narrative environment and publicly available to users.

**Figure 2.**
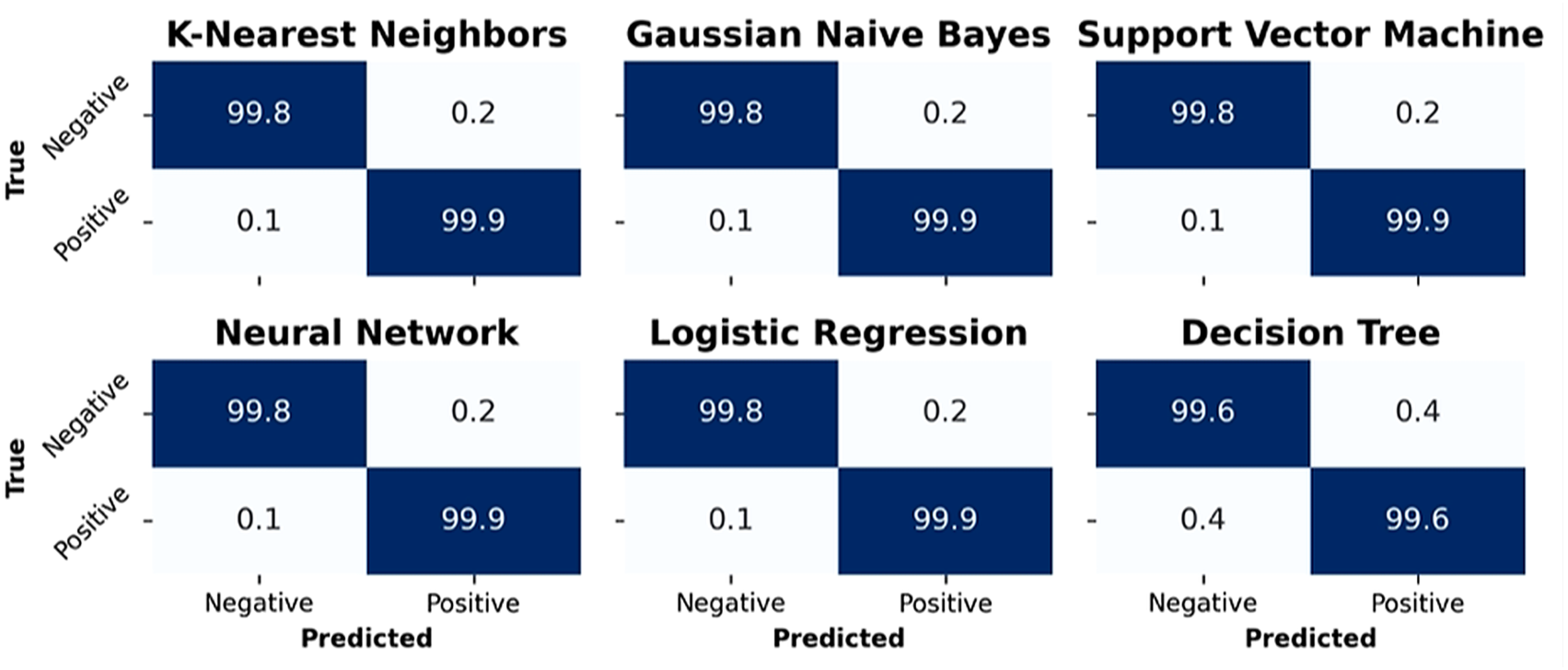
Prediction of Gram-stain type using six machine learning classifiers. Confusion matrices show strong performance across classifiers, including the tuned decision tree, in distinguishing Gram-negative and Gram-positive bacteria.

**Figure 3.**
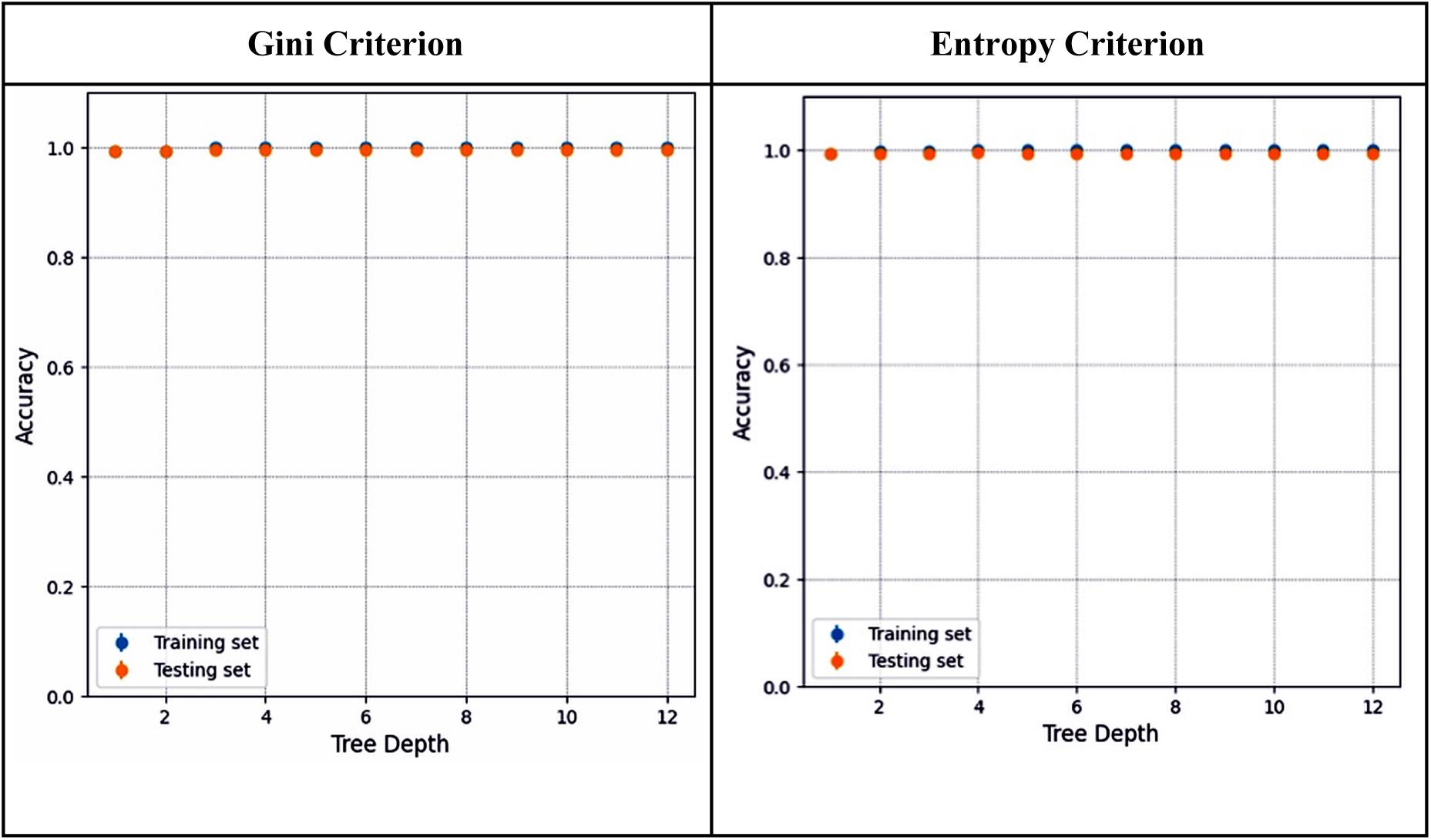
Comparison of Accuracy between Gram stain training versus testing sets based on the Gini Criterion and the Entropy Criterion for 12 levels of decision tree depth indicates the Entropy Criterion and a tree depth of 4 are the optimal hyperparameters.

**Figure 4.**
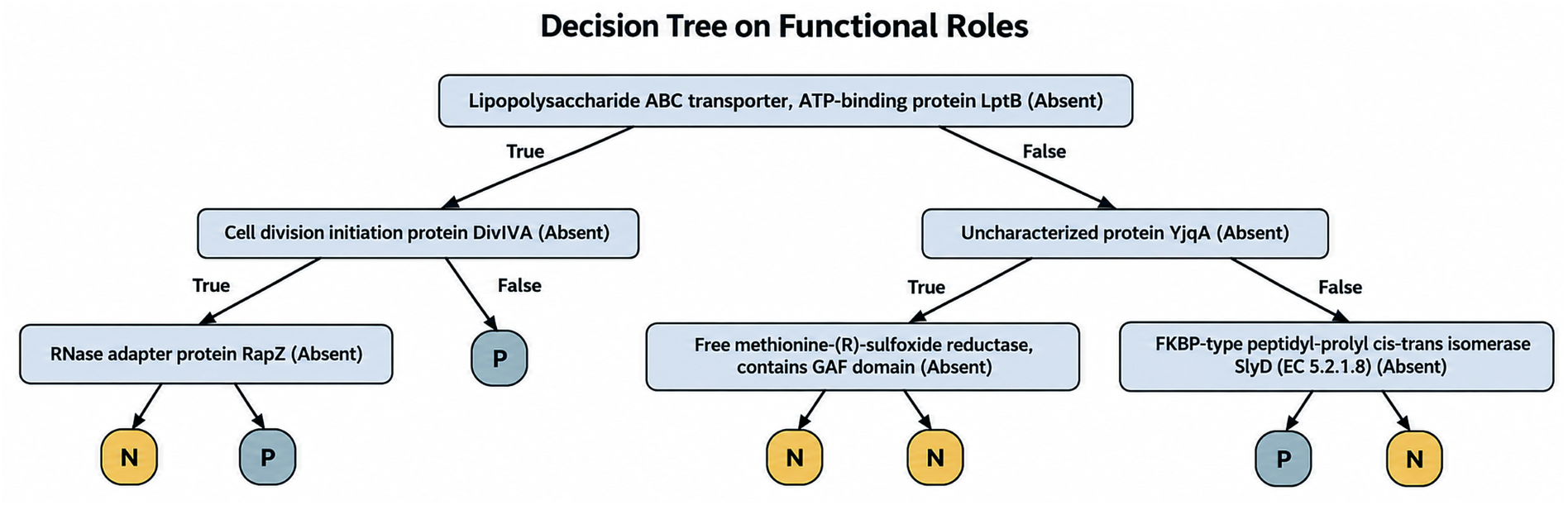
Decision tree illustrating an interpretable pathway for classifying bacterial Gram type based on RAST-annotated functional roles. Each decision node evaluates the presence or absence of a specific functional role, and the terminal leaf nodes indicate the predicted class as either Gram-positive (P) or Gram-negative (N).

## Results

The performance of all the classifiers was evaluated by comparing the confusion matrix of each classifier along with the performance metrics: accuracy, precision, recall, and F1 Score. Furthermore, the classifier performance was tested against Gram type and respiration type separately. Finally, the best performing decision tree classifiers for the Gram, the respiration and the growth phenotypes were analyzed to enable biological interpretation of the misclassified genomes.

### Gram Classifiers

The confusion matrices illustrated in Figure 2 show the average Gram stain prediction scores across ten cross-validation testing folds. Each value is a percentage of the total number of the true class. On average, each test set consists of 90 Gram-negative and 135 Gram-positive bacteria (10% of the total samples in each category). We observe that the classifiers predict the held-out genomes with near perfect accuracy, noting that the tuned decision tree classifier has slightly lower performance. Further analysis of the confusion matrices reveals that the precision and recall are both >0.99 for all the classifiers, indicating that each of the models has strong predictive power on the Gram stain task. Given the interpretability of the decision tree, we tuned its accuracy with respect to two parameters: the split quality criterion (Gini impurity or Entropy) and the tree depth (see Methods: Decision Tree Tuning). As depicted in Figure 3, each criterion was evaluated against 12 levels of tree depth by comparing the accuracy on the training and testing sets. We observe that the decision tree is a robust predictor for Gram stain, predicting with near-perfect accuracy (precision and recall >0.99) on both the training and testing sets across all of the hyperparameter combinations, with the optimal tree using Entropy criterion and a tree depth of 4.

### Biological Interpretation

Using the optimized decision tree classifier hyperparameters (Entropy criterion and a tree depth of 4), the full classification tree orders the functional roles by importance for the Gram stain prediction task. As illustrated in Figure 4, the decision tree bases its classification on the presence or absence of different functional roles (purple) with the final leaf nodes indicating the Gram stain as positive (P) or negative (N). We further enumerate the significant functional roles and respective feature importance in Supp. Table 1.

Interpreting the decision tree classifier provides a biologically meaningful method for distinguishing between Gram-negative and Gram-positive given the functional roles. In Gram-positive bacteria, the presence of the cell division initiation protein DivIVA is unique and plays a crucial role in the cell division process (Thomaides et al. 2001). On the other hand, in Gram-negative bacteria, the Lipopolysaccharide ABC transporter, ATP-binding protein LptB, is associated with drug resistance and functions as an inter-membrane protein (Dong et al. 2017). Another notable functional role specific to Gram-negative bacteria is the free methionine-(R)-sulfoxide reductase-GAF domain protein, which has been extensively characterized in Escherichia coli (Lin et al. 2007). These distinct functional roles serve as key markers for differentiating between Gram-negative and Gram-positive bacteria, contributing to our understanding of their unique characteristics and biological processes.

### Respiration Classifiers

Predicting respiration type is a more challenging task than predicting Gram stain, in part, due to the presence of 3 class labels (aerobe, anaerobe, and facultative). The confusion matrices shown in Figure 5 depict the average classifier performance on the testing sets generated via ten-fold cross-validation for the task of predicting respiration type given the RAST functional role attributes. Each value is a percentage of the total number of the true class, where the validation sets consist, on average, of 25 aerobe, 13 anaerobe, and 17 facultative types (10% of the total samples in each category). As in the Gram stain case study (See Results: Gram Classifiers), we report the analysis based on a tuned decision tree (Entropy criterion, tree depth 8). As illustrated in Figure 6, we observe that as the depth of the decision tree increases, it is able to express more complex relationships within the training data, however, increasing tree depth can also lead to overfitting and poorer generalization to held-out data. We select the optimal hyperparameters by selecting the maximum performance on the testing set, just before overfitting occurs. Further analysis of the confusion matrices as listed in Table 4, indicates that the Gaussian Naive Bayes classifier outperformed the rest of the classifiers with an overall prediction accuracy of 81.5%. We note that the tuned decision tree (accuracy of 79.6%) is competitive with several other models, namely, K-Nearest Neighbors, Neural Network, and Logistic Regression, and also outperforms the Support Vector Machine and the untuned decision tree, which both have an accuracy of 77.8%.

**Figure 5.**
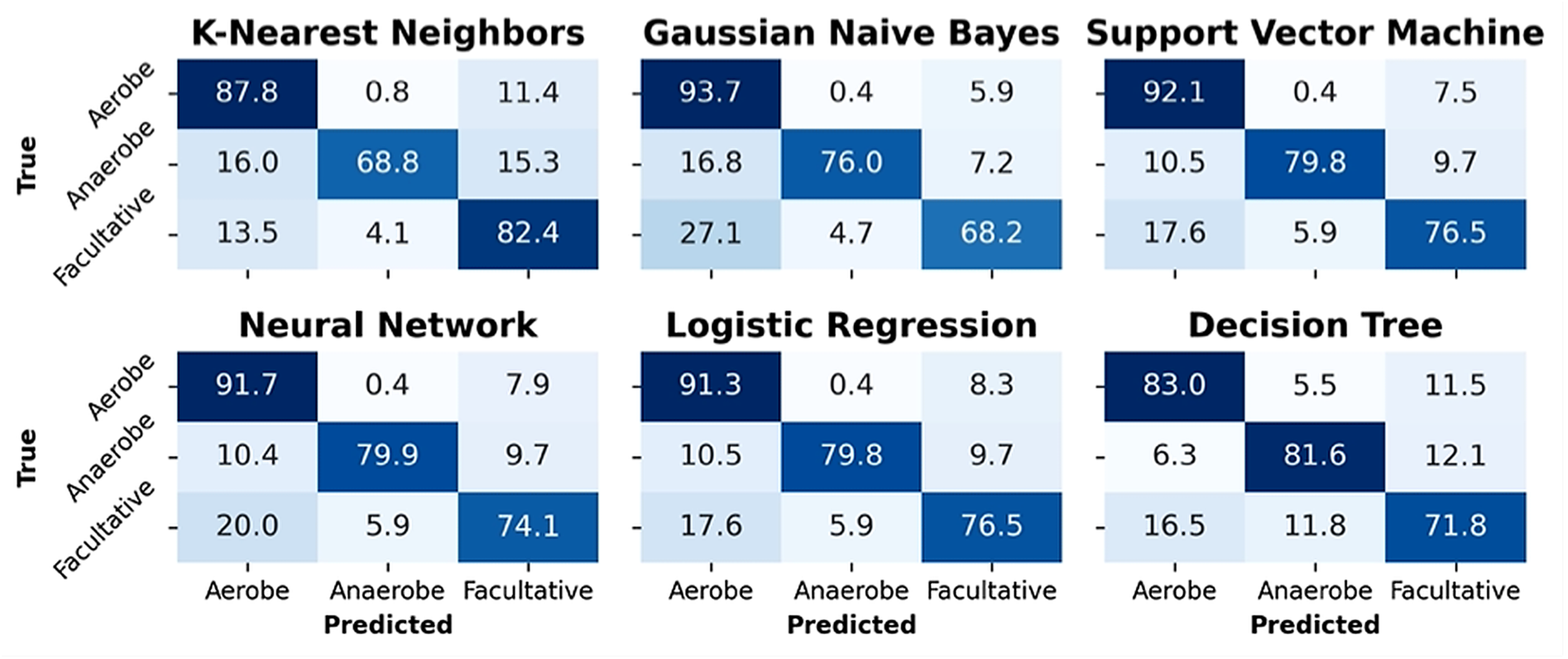
Prediction performance of respiration type (aerobe, anaerobe, and facultative) based on six classifiers including a tuned decision tree (Entropy criterion, tree depth 8).

**Figure 6.**
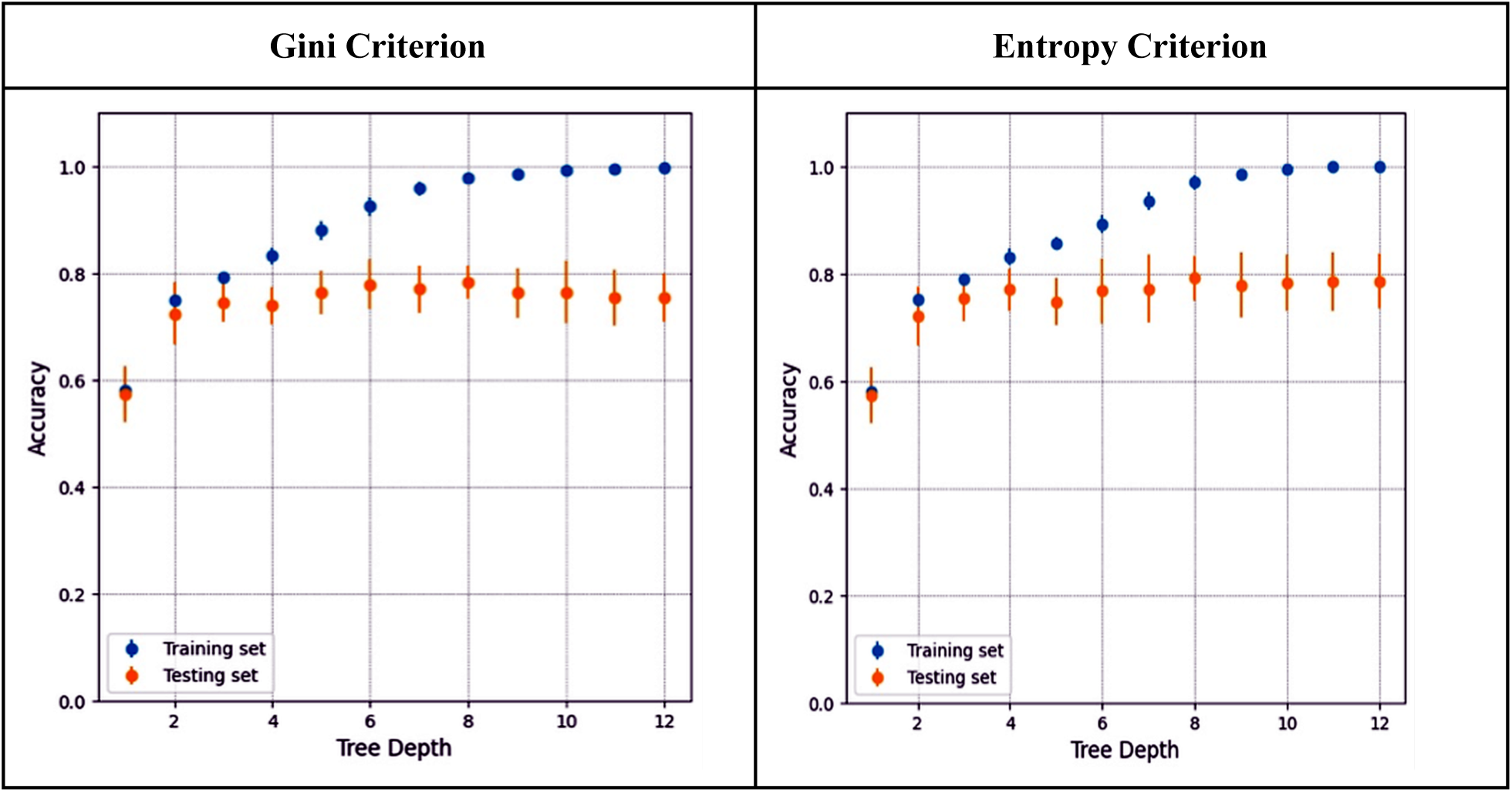
Hyperparameter tuning of the decision tree criterion and tree depth evaluated via accuracy on the training and testing sets for the task of predicting respiration type indicates the optimal parameters are the Entropy criterion with a tree depth of 8.

**Table 4.** Model performance statistics in the form of Accuracy, Precision, Recall, and F1 Score calculated against the confusion matrices of respiration type for the six classifiers and a tuned decision tree (Entropy criterion, tree depth 8).

|  | K-Nearest<br>Neighbors | Gaussian<br>Naive Bayes | Logistic<br>Regression | Untuned<br>Decision<br>Tree | Tuned<br>Decision<br>Tree | Support<br>Vector<br>Machine | Neural<br>Network |
| --- | --- | --- | --- | --- | --- | --- | --- |
| **Aerobe** |  |  |  |  |  |  |  |
| Precision | 0.759 | 0.786 | 0.808 | 0.8 | 0.870 | 0.778 | 0.808 |
| Recall | 0.88 | 0.88 | 0.84 | 0.8 | 0.8 | 0.84 | 0.84 |
| F1 score | 0.815 | 0.830 | 0.824 | 0.8 | 0.833 | 0.808 | 0.824 |
| **Anaerobe** |  |  |  |  |  |  |  |
| Precision | 1 | 0.909 | 0.9 | 0.9 | 0.833 | 0.9 | 0.909 |
| Recall | 0.75 | 0.833 | 0.75 | 0.75 | 0.833 | 0.75 | 0.833 |
| F1 score | 0.857 | 0.870 | 0.818 | 0.818 | 0.833 | 0.818 | 0.870 |
| **Facultative** |  |  |  |  |  |  |  |
| Precision | 0.75 | 0.8 | 0.722 | 0.684 | 0.684 | 0.706 | 0.706 |
| Recall | 0.706 | 0.706 | 0.765 | 0.765 | 0.765 | 0.706 | 0.706 |
| F1 score | 0.727 | 0.75 | 0.743 | 0.722 | 0.722 | 0.706 | 0.706 |
| **Accuracy** | 0.796 | **0.815** | 0.796 | 0.778 | **0.796** | 0.778 | 0.796 |

### Biological Interpretation

Unlike the Gram stain phenotype, the optimized decision tree (Entropy criterion and a tree depth of 8) for the respiration task is too large to report as a figure. Instead, we enumerate the significant functional roles and their respective feature importance in Supp. Table 2 and link to the full report in KBase with the plotted decision tree in Appendix A2.

Classification of organisms based on respiration type involves the identification of specific functional roles associated with aerobic and anaerobic metabolism. In strict anaerobes such as Clostridium, the gene encoding the NADH-dependent reduced ferredoxin:NADP+ oxidoreductase subunit A has been identified (Liang et al. 2019) with a high importance score. This gene is commonly found in the cytoplasm of various anaerobic bacteria and archaea (Demmer et al. 2015). It serves as a reliable differentiator for classifying aerobic and anaerobic organisms.

Another crucial functional role for distinguishing anaerobes and facultative anaerobes from aerobes is the anaerobic enzyme Ribonucleotide reductase of class III (anaerobic), large subunit (EC 1.17.4.2), which is present in anaerobes and facultative anaerobes like Escherichia coli (Eliasson et al. 1990). This enzyme is prominently ranked among the functional roles that differentiate anaerobic and facultative anaerobic organisms from aerobes. Furthermore, the role of ’Flavocytochrome c:sulfide dehydrogenase’ also shows a relatively high score in our decision tree classifier, indicating its importance in determining respiration type. This role encodes the enzyme responsible for catalyzing the oxidation of sulfide and polysulfide ions to elemental sulfur to cytochrome c in haloalkaliphilic obligate autotroph anaerobes (Tikhonova et al. 2021) and is also present in the facultative aerobic organism *Paracoccus denitrificans* (Kostanjevecki et al. 2000). Finally, the activator protein of ’(R)-2-hydroxyglutaryl-CoA dehydratase’ is another key feature that tends to be present in anaerobes. This enzyme, isolated from *Acidaminococcus fermentans* under strict anaerobic conditions, plays a role in the fermentation of glutamate via the hydroxyglutarate pathway, producing ammonia, carbon dioxide, acetate, butyrate, and molecular hydrogen (Muller and Buckel 1995). In summary, these specific functional roles provide valuable insights into classifying organisms based on their respiration types, distinguishing between aerobic, anaerobic, and facultative organisms.

### Phenotype Growth Study Classifiers

The confusion matrices illustrated in Figure 7 show the average phenotype growth prediction scores across ten cross-validation testing folds for N-Acetyl-D-Glucosamine substrate. Each value represents the percentage of samples in the corresponding true class, where 0 denotes no growth and 1 denotes growth. We used 10-fold stratified cross-validation (K = 10); thus, each test fold contained approximately 14 genomes, including about 9 growth and 5 no-growth phenotypes.

Each value represents the percentage of samples in the corresponding true class, where 0 denotes no growth and 1 denotes growth. We used 10-fold stratified cross-validation (K = 10); thus, each test fold contained approximately 14 genomes, including about 9 growth and 5 no-growth phenotypes.

**Figure 7.**
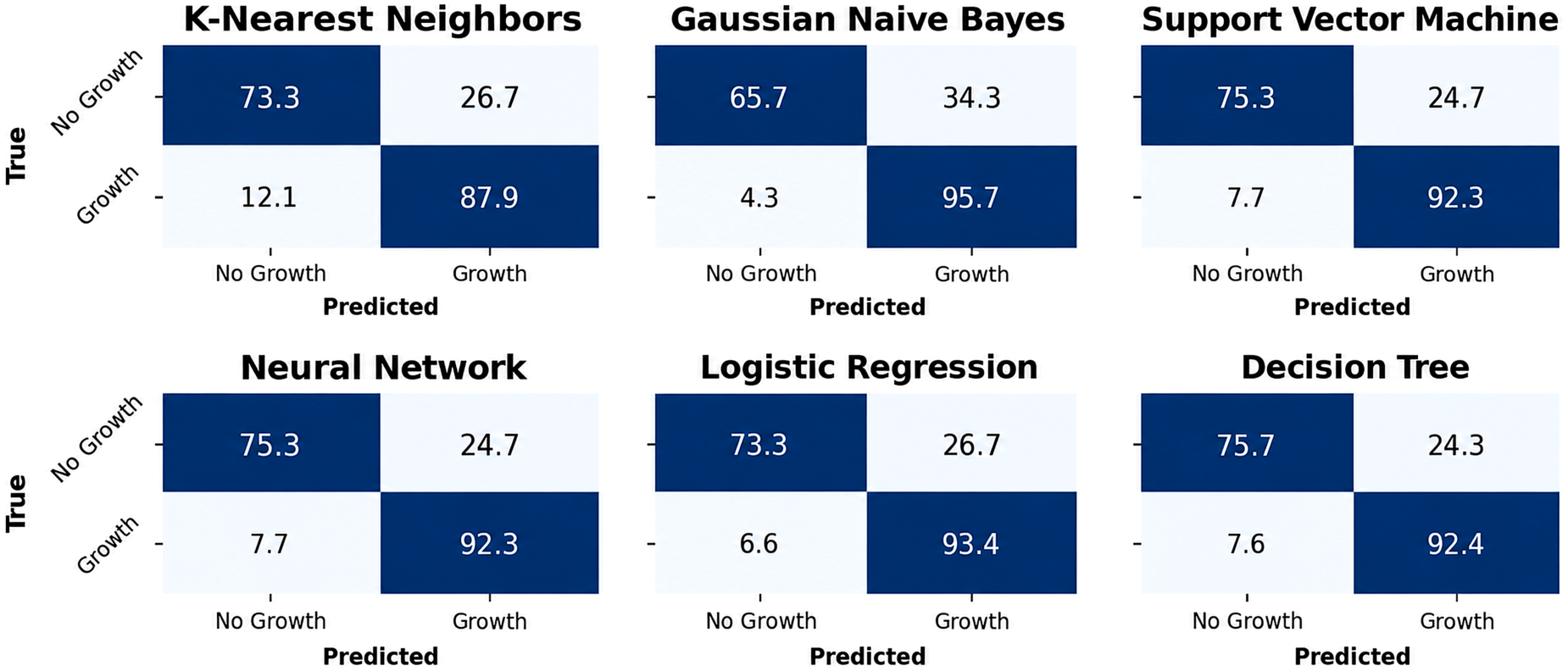
Prediction performance of growth type (growth or no growth) based on six classifiers including a tuned decision tree (Entropy criterion, tree depth 8) for one of the 64 substrates, namely, N-Acetyl-D-Glucosamine.

For the held-out phenotypes comprising the test set, as illustrated in Figure 7, all classifiers predict the phenotype growth types when there is growth with minimum accuracy of 87.9% for K-nearest Neighbors and maximum accuracy of 95.7% for Gaussian Naive Bayes. In cases where there is no growth, the minimum accuracy was found to be 65.7% with the gaussian naive bayes and the maximum accuracy was found to be 75.7% with the decision tree. Similar to the Gram stain and Respiration case studies, we tuned the decision tree with the split quality criterion (Gini impurity or Entropy) and the tree depth. As depicted in Figure 8, the tuning process was the same as the other two case studies i.e. each criterion being evaluated against 12 tree depth levels. Both gini impurity and entropy tuned decision trees performed remarkably well when there is growth as seen by their 94.6% and 95.7% accuracy respectively. In cases where there is no growth, gini impurity and entropy tuned decision trees achieved 81.3% and 83.3% accuracy respectively, compared to the untuned decision tree which showed 75.7% accuracy.

**Figure 8.**
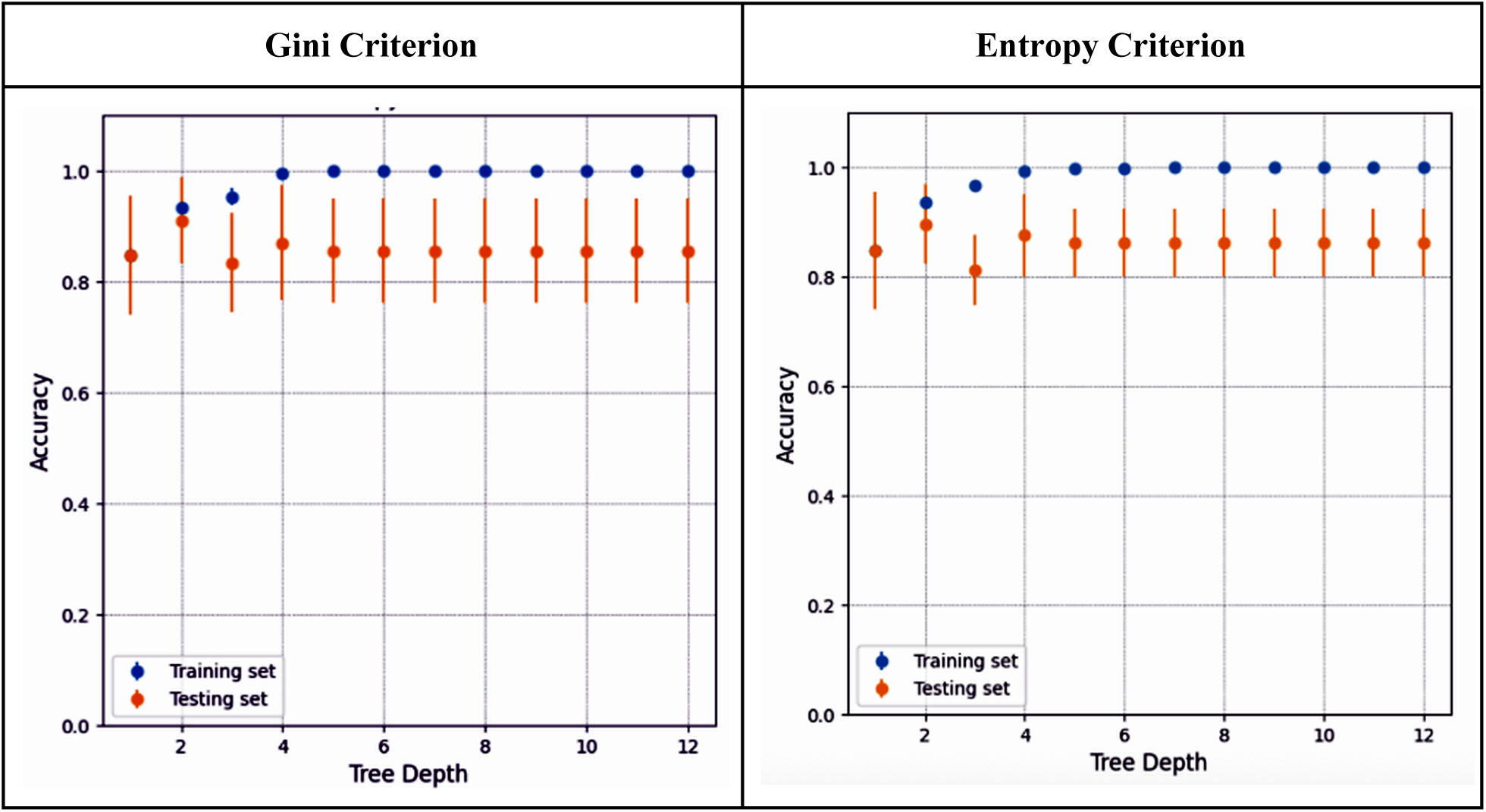
Hyperparameter tuning of the decision tree criterion and tree depth evaluated via accuracy on the training and testing sets for the task of predicting growth type indicates the optimal parameters are the Entropy criterion with a tree depth of 8 for N-Acetyl-D-Glucosamine substrate.

Table 5 shows the precision, recall, F1 score and accuracy for all classifiers, including the tuned decision trees for N-Acetyl-D-Glucosamine substrate. The tuned decision trees showed performance comparable to that of the untuned decision tree across precision, recall, and F1 score for both growth classes (label 1, growth; label 0, no growth). In contrast, the neural network achieved the highest overall accuracy, with an accuracy of approximately 0.86. Gaussian Naive Bayes underperformed with ∼0.71 accuracy compared to the rest of the five classifiers. Interestingly, gini impurity tuned and entropy tuned decision trees did not improve the performance relative to the untuned decision tree (see Figure 9).

**Figure 9.**
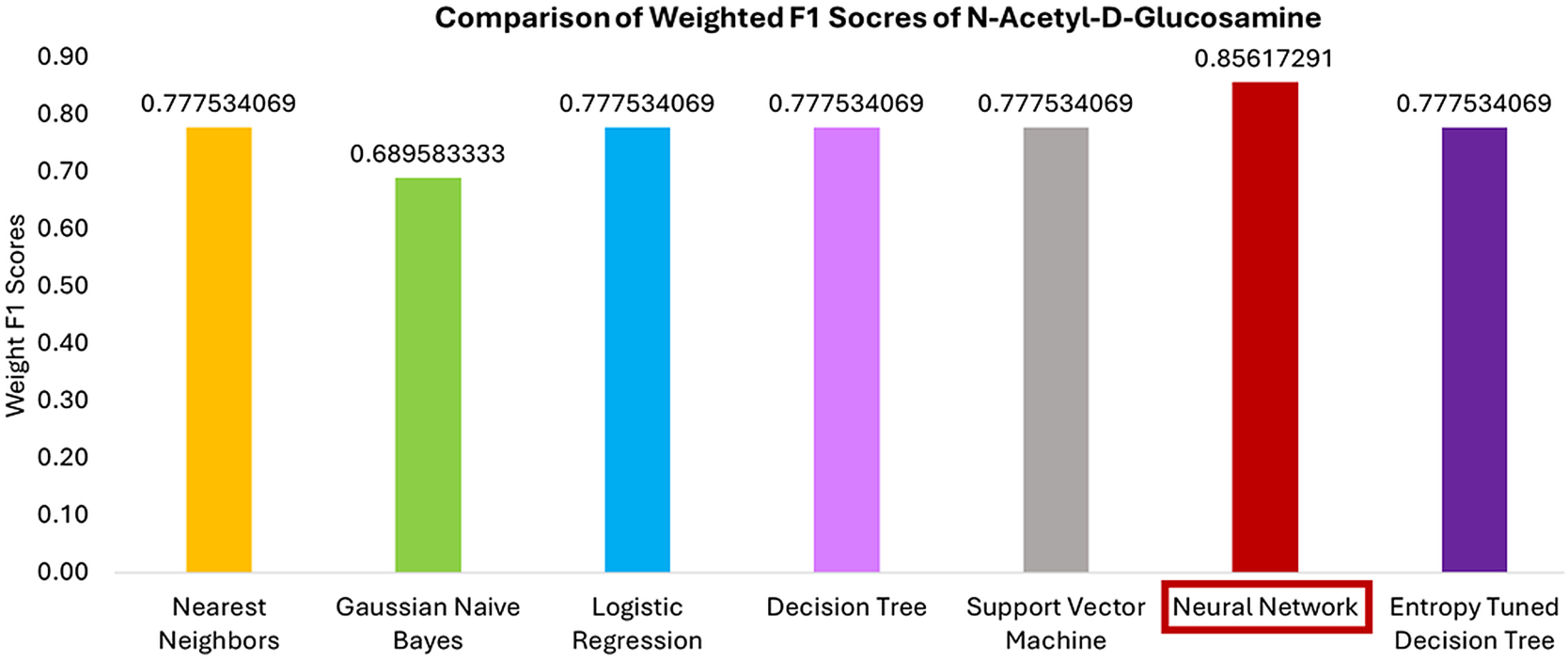
Weighted F1 scores of all six classifiers for growth or no growth phenotype prediction in N-Acetyl-D-Glucosamine substrate.

**Table 5.** Model performance statistics in the form of Accuracy, Precision, Recall, and F1 Score calculated against the confusion matrices of growth type for the six classifiers and a tuned decision tree (Entropy criterion, tree depth 8) for N-Acetyl-D-Glucosamine substrate.

| Metrics | K Nearest Neighbors | Gaussian Naive Bayes | Logistic Regression | Decision Tree | Decision Tree Gini | Decision Tree Entropy | Support Vector Machine | Neural Network |
| --- | --- | --- | --- | --- | --- | --- | --- | --- |
| 0 |  |  |  |  |  |  |  |  |
| Precision | 0.75 | 0.666667 | 0.75 | 0.75 | 0.75 | 0.75 | 0.75 | 0.8 |
| Recall | 0.6 | 0.4 | 0.6 | 0.6 | 0.6 | 0.6 | 0.6 | 0.8 |
| F1-Score | 0.666667 | 0.5 | 0.666667 | 0.666667 | 0.666667 | 0.666667 | 0.666667 | 0.8 |
| 1 |  |  |  |  |  |  |  |  |
| Precision | 0.8 | 0.727273 | 0.8 | 0.8 | 0.8 | 0.8 | 0.8 | 0.888889 |
| Recall | 0.888889 | 0.888889 | 0.888889 | 0.888889 | 0.888889 | 0.888889 | 0.888889 | 0.888889 |
| F1-Score | 0.842105 | 0.8 | 0.842105 | 0.842105 | 0.842105 | 0.842105 | 0.842105 | 0.888889 |
| Accuracy | 0.785714 | 0.7142286 | 0.785714 | 0.785714 | 0.785714 | 0.785714 | 0.785714 | 0.857143 |

### Biological Interpretation

Decision trees provide interpretable and biologically meaningful insights into the functional roles that differentiate phenotypes exhibiting growth from those showing no growth on specific substrates. Figures 10-A and 10-B illustrate the decision trees identifying key functional roles associated with N-Acetyl-D-Glucosamine and quinic acid utilization, respectively.

**Figure 10.**
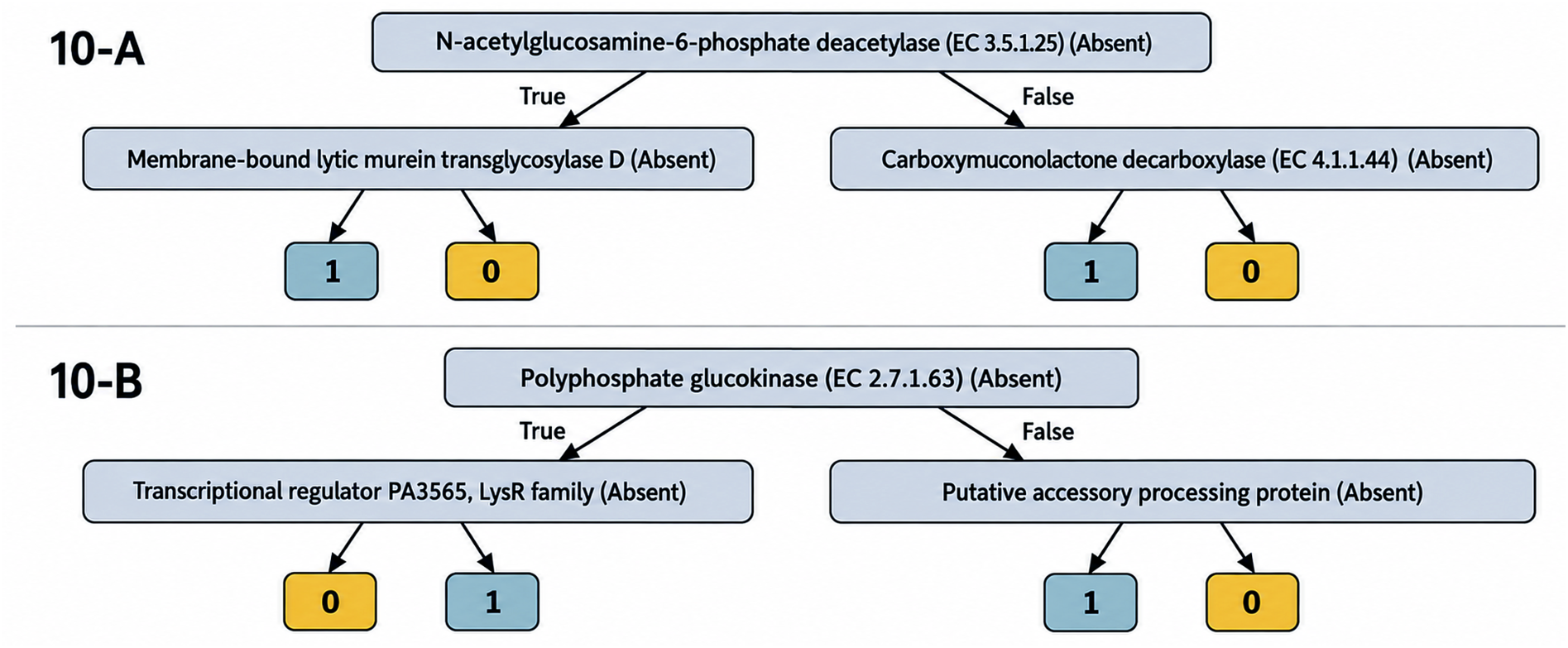
Decision trees illustrating interpretable predictive pathways for bacterial growth phenotypes based on RAST-annotated functional roles. Each decision node evaluates the presence or absence of a specific functional role, and the terminal leaf nodes indicate the predicted phenotype as either growth (1) or no growth (0). (A) Decision tree for growth on N-acetyl-D-glucosamine. (B) Decision tree for growth on Quinic acid.

N-acetylglucosamine-6-phosphate deacetylase (NagA) emerged as an informative feature. NagA is involved in the amino sugar metabolism pathway that converts N-acetylglucosamine-6-phosphate (GlcNAc-6-P) to glucosamine-6-phosphate (GlcN-6-P), and is part of the conserved *nag* regulon in bacteria (Plumbridge 1990, 1991; Yadav et al. 2011). If NagA is selected as the root node in the decision tree, this suggests that the capacity to metabolize amino sugars is discriminative between growth phenotypes. The membrane-bound lytic murein transglycosylase D (MltD) plays an important role during cell growth and division, potentially influencing fitness under different nutrient conditions (UniProt Consortium 2026). On the right-side branch, carboxymuconolactone decarboxylase is involved in the beta-ketoadipate pathway which degrades aromatic compounds and reflects a broad metabolic versatility (Harwood and Parales 1996; Pérez-Pantoja et al. 2008). The decision tree selecting these functional roles suggests that metabolic breadth and cell envelope physiology are correlated with the observed growth phenotypes, rather than direct catabolism of the tested substrate. For Quinic-Acid, polyphosphate glucokinase (EC 2.7.1.63) enzyme was found to be the most important predictor. This enzyme phosphorylates glucose to glucose-6-phosphate using inorganic polyphosphate or polyP instead of ATP and thus presents an alternative pathway to energy metabolism (Tanaka et al. 2003). Regardless of the substrate, tuned decision trees were broadly useful in identifying the functional roles whose presence or absence helps distinguish between phenotypes exhibiting growth or no growth.

## Discussion and Conclusion

In this work we demonstrate how ML can be combined with RAST functional annotation to automatically classify bacterial genome phenotypes. We have validated the approach using six ML algorithms (K-Nearest Neighbors, Gaussian Naive Bayes, Support Vector Machines, Neural Networks, Logistic Regression, and Decision Trees) against a high-quality literature-derived training set in order to predict three selected bacterial phenotypes: (i) Gram stain; (ii) Respiration type of aerobic, anaerobic, and facultative; and (iii) growth or no-growth across a diverse set of carbon sources. In our analysis, we show that for the Gram stain prediction task, all the classifiers can predict the correct phenotype with near-perfect accuracy (both precision and recall >0.99). On the more challenging task of predicting the respiration type, we achieve an overall prediction accuracy across aerobic, anaerobic, and facultative of ∼81.5% using a Gaussian Naive Bayes classifier. For predicting growth versus no-growth phenotypes, we achieved an accuracy of ∼86% using a neural network. This level of performance is especially noteworthy given the limited size of the training dataset available for each carbon source. We anticipate that as the dataset expands to include a larger and more diverse set of organisms, the predictive accuracy of the model will further improve.

We also demonstrate how tuned decision tree classifiers can be interpreted to reveal biological insights implicating the presence (or absence) of key functional roles that are related to the biochemical role of the associated phenotype, thus helping biologists understand the predicted phenotype (see Results). The complete phenotype prediction pipeline, comprising four applications, is readily accessible through The Department of Energy Systems Biology Knowledgebase (KBase) (Appendix A2) enabling users to process their training-set data, build the six classifiers described here, and predict phenotypes for unclassified genomes in a high-throughput manner. Furthermore, users can take advantage of this open access platform to access the datasets used in this study and effortlessly reproduce the results that are discussed in this study.

This study also paves the way to future research that would involve identifying genomes that do not fit well into established classes and exploring the possibility of defining new potential classes. For example, some organisms may not fit conventional Gram-positive or Gram-negative categories, thereby challenging binary Gram-stain classification. This highlights the potential of ML to not only classify well-known classes but also uncover and characterize new classes. Such investigations could contribute significantly to expanding our understanding of microbial diversity and show the utility of ML in capturing complex biological variations.

ML classifiers can also be utilized to identify and explore phenotypes that are not well explained in terms of enzyme function or have yet to be fully characterized in metabolic pathways. One of the clear advantages of ML classifiers is their ability to capture the comprehensive functional attributes of organisms. While enzymatic functions related to specific phenotypes may be absent, this holistic approach enables ML classifiers to predict these phenotypes with reasonable accuracy, surpassing the limitations of relying solely on enzyme function annotation algorithms or biochemical pathways. By coupling ML predicted phenotypes with methods such as mechanistic modeling, we can uncover novel associations and gain a deeper understanding of complex phenotypes in a more comprehensive and efficient manner.

Finally, while this study focused solely on genome annotations to build predictor attributes for classification algorithms, the approach could be extended to use protein sequences, functional domains, short DNA sequences, or biochemical reactions as attributes. In summary, our technique establishes an accurate and efficient method to predict important bacterial phenotypes that, we believe, can be applied to predict phenotypes such as antimicrobial resistance or pathogenicity that are medically and industrially important.

## Appendix

## A1 Machine Learning Software

Scikit-learn (version 0.23.2) default parameters used in this study.

~~~
sklearn.neighbors.**KNeighborsClassifier**(n_neighbors=5, *, weights=’uniform’, algorithm=’auto’, leaf_size=30, p=2, metric=’minkowski’, metric_params=None,
n_jobs=None)
sklearn.linear_model.**LogisticRegression**(*penalty=’l2’*, *\**, *dual=False*, *tol=0.0001*, *C=1.0*, *fit_intercept=True*, *intercept_scaling=1*, *class_weight=None*, *random_state=None/0*, *solver=’lbfgs’*, *max_iter=100*, *multi_class=’auto’*, *verbose=0*, *warm_start=False*, *n_jobs=None*, *l1_ratio=None*)
sklearn.naive_bayes.**GaussianNB**(*\**, *priors=None*, *var_smoothing=1e-09*)
sklearn.svm.**SVC**(*\**, *C=1.0*, *kernel=’rbf’*, *degree=3*, *gamma=’scale’*, *coef0=0.0*, *shrinking=True*, *probability=False*, *tol=0.001*, *cache_size=200*, *class_weight=None*, *verbose=False*, *max_iter=-1*, *decision_function_shape=’ovr’*, *break_ties=False*, *random_state=None*)
sklearn.neural_network.**MLPClassifier**(*hidden_layer_sizes=(100,)*, activation*=’relu’*, *\**, *solver=’adam’*, *alpha=0.0001*, *batch_size=’auto’*, *learning_rate=’constant’*, *learning_rate_init=0.001*, *power_t=0.5*, *max_iter=200*,
~~~

*shuffie=True, random_state=None, tol=0.0001, verbose=False, warm_start=False, momentum=0.9, nesterovs_momentum=True, early_stopping=False*,

*validation_fraction=0.1*, *beta_1=0.9*, *beta_2=0.999*, *epsilon=1e-08*, *n_iter_no_change=10*, *max_fun=15000*)

~~~
sklearn.tree.**DecisionTreeClassifier**(*\**, *criterion=’gini’*, *splitter=’best’*, *max_depth=None*, *min_samples_split=2*, *min_samples_leaf=1*, *min_weight_fraction_leaf=0.0*, *max_features=None*, *random_state=None*,
~~~

*max_leaf_nodes=None, min_impurity_decrease=0.0, class_weight=None, ccp_alpha=0.0)*

### Supplemental Tables

**Supp. Table 1.** Functional roles used in the Gram Decision Tree classifier and the respective feature importance.

| Functional Role | Importance |
| --- | --- |
| Lipopolysaccharide ABC transporter, ATP-binding protein LptB | 9.358330e-01 |
| Cell division initiation protein DivIVA | 3.866963e-02 |
| RNase adapter protein RapZ | 1.335353e-02 |
| Uncharacterized protein YjqA | 8.269172e-03 |
| FKBP-type peptidyl-prolyl cis-trans isomerase SlyD (EC 5.2.1.8) | 3.874681e-03 |
| Free methionine-(R)-sulfoxide reductase, contains GAF domain | 6.574823e-18 |

**Supp. Table 2.**
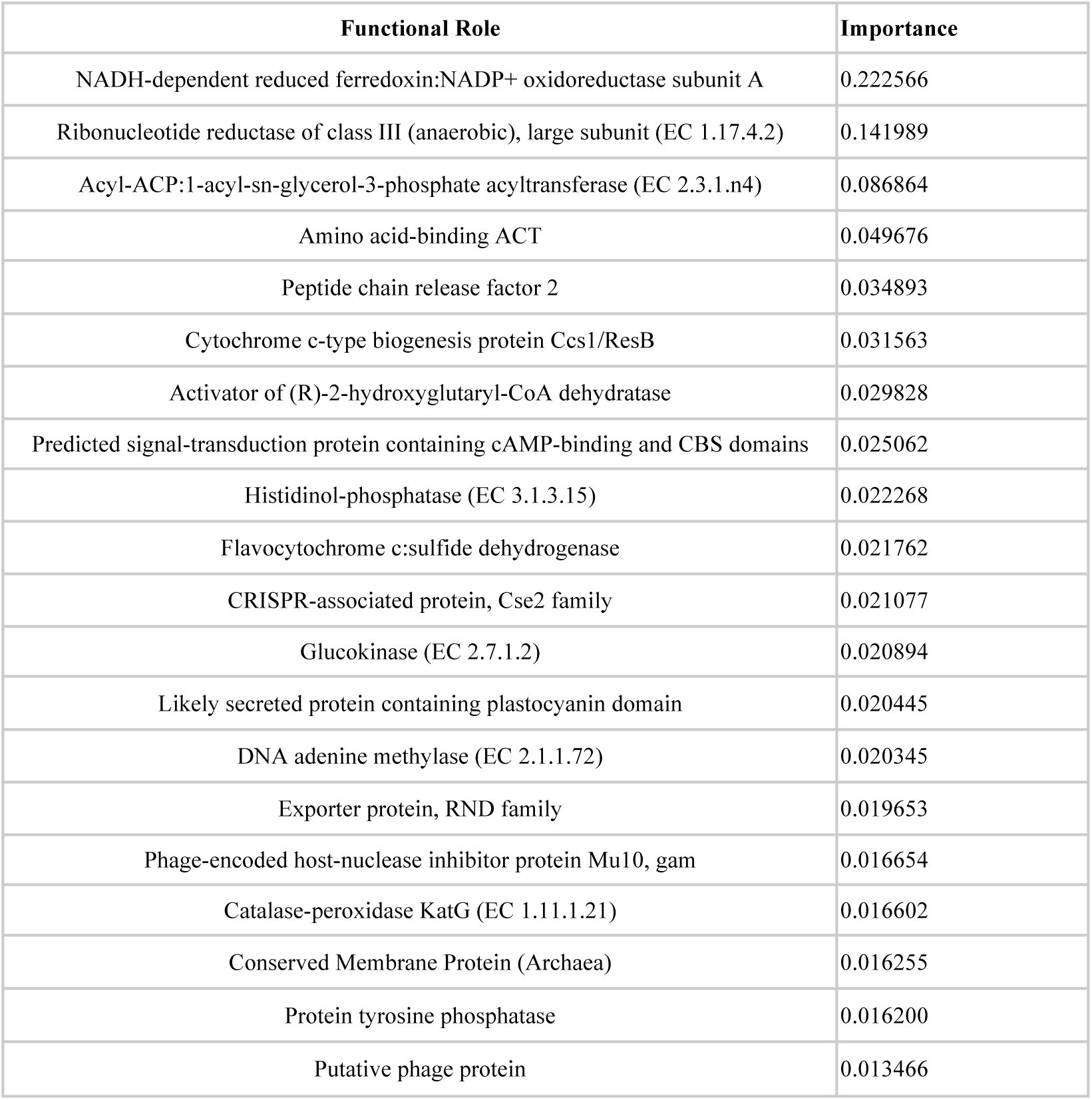
Functional roles used in the Respiration Decision Tree classifier and the respective feature importance.

**Supp. Table 3.**
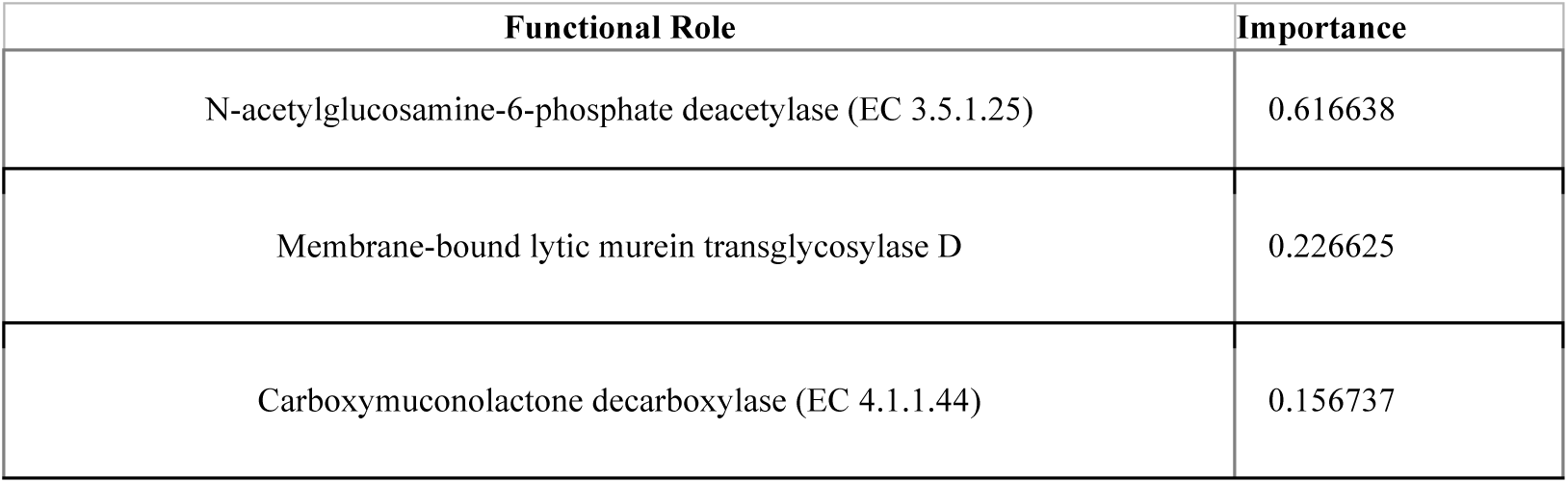
Functional roles used in the growth type Decision Tree classifier and the respective feature importance for importance for N-Acetyl-D-Glucosamine substrate.

| Functional Role | Importance |
| --- | --- |
| N-acetylglucosamine-6-phosphate deacetylase (EC 3.5.1.25) | 0.616638 |
| Membrane-bound lytic murein transglycosylase D | 0.226625 |
| Carboxymuconolactone decarboxylase (EC 4.1.1.44) | 0.156737 |

## A2 Source Code, KBase Narratives and Applications

Source code: https://github.com/janakagithub/kb_genomeclassification

Gram case study KBase narrative: https://narrative.kbase.us/narrative/153575

Respiration case study KBase narrative: https://narrative.kbase.us/narrative/153678

Phenotype growth case study KBase narrative: https://narrative.kbase.us/narrative/137002

KBase modules designed for this study: https://narrative.kbase.us/#catalog/modules/kb_genomeclassification

The KBase pipeline is comprised of four applications (i) Upload high-quality data to train classifiers; (ii) Annotate genomes in the training set with the RAST annotation algorithm; (iii) Build six different genome classifiers; and (iv) Predict the phenotype of unclassified genomes. Screenshots for each app and descriptions are provided below.

(i) Upload high-quality data to train classifiers:

**Supp. Figure 1.**
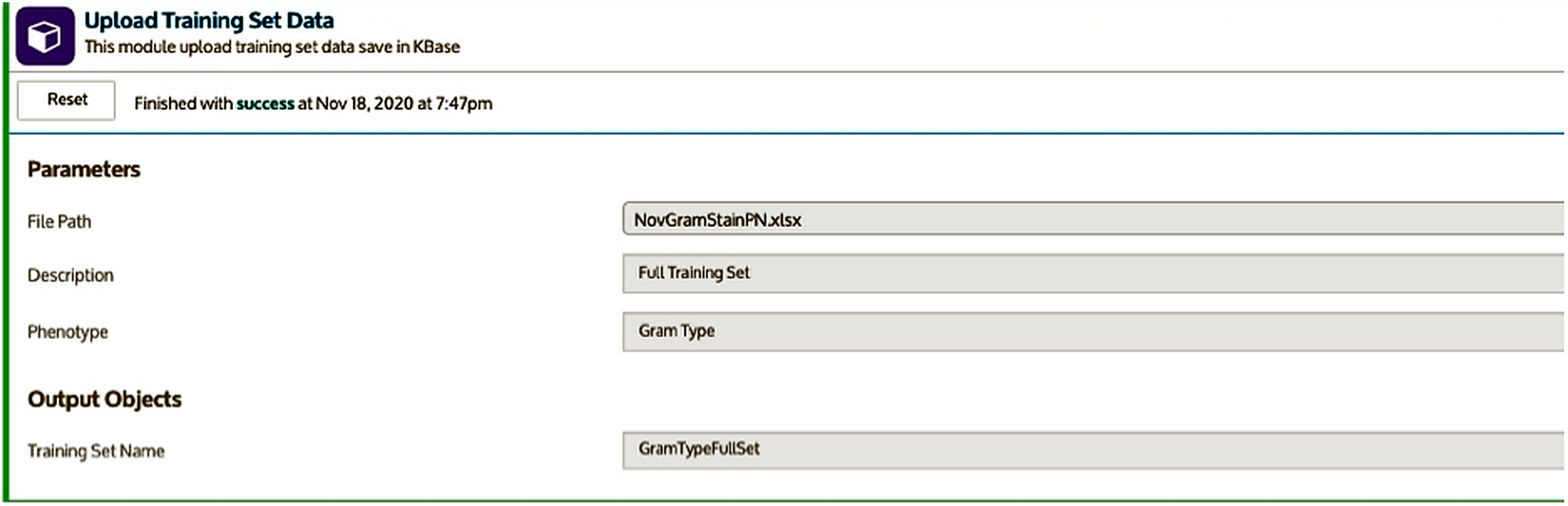
*Upload Training Set Data* app. This app takes a training set that comprises genomes and respective phenotypes. At minimum, the input file is required to contain two columns: genome name/ID and phenotype. This app reads the genomes and phenotype data then builds a TrainingSet data object associating the phenotypes to respective genomes.

Template for Upload Training Set Data App

Required - 2 columns: (Genome Name || Genome Reference || Ref Seq Ids) && Phenotype Optional (semicolon separated list): References, Evidence Types

Example:

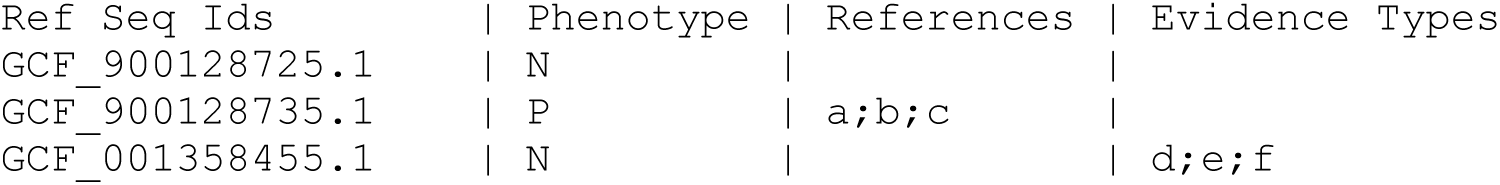

(ii) Annotate genomes in the training set with the RAST annotation algorithm:

**Supp. Figure 2.**
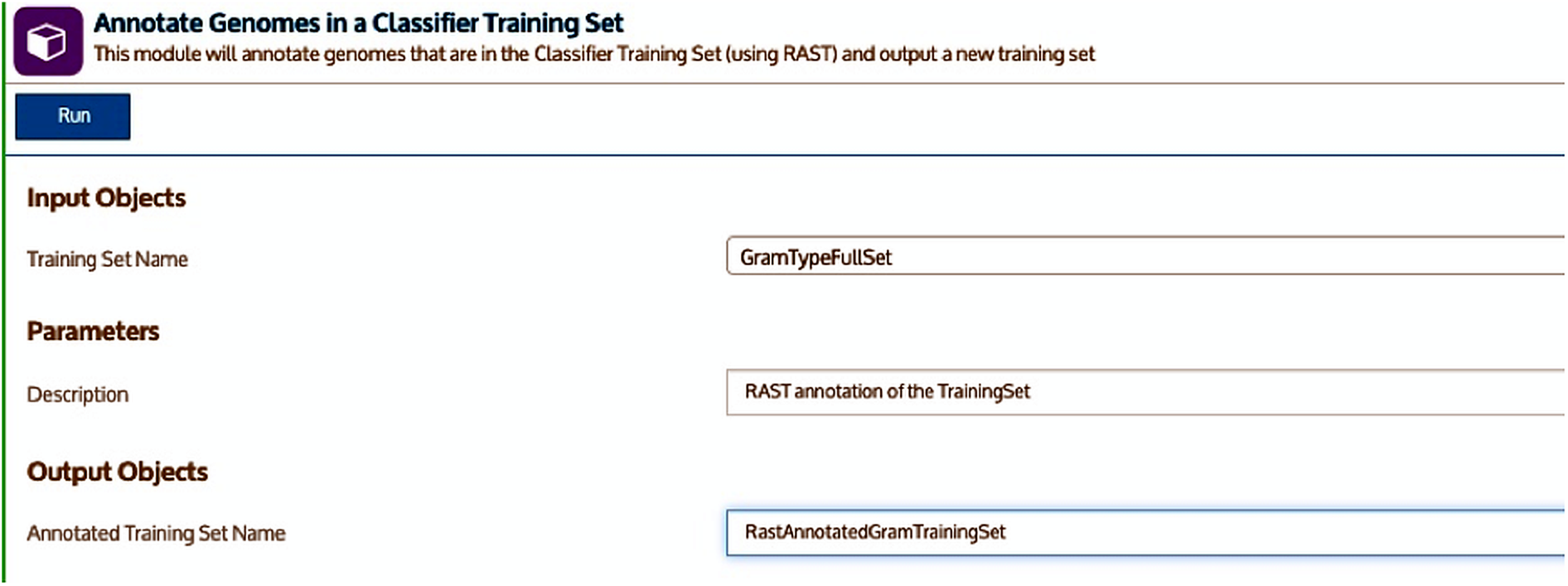
*Annotate Genomes in a Classifier Training Set* app. This app uses the training set that was generated from the *Upload Training Set Data* app and annotates all genomes in the training set using the RAST annotation algorithm. The RAST annotations are being used as predictors for constructing classifiers.
(iii) Build six different genome classifiers:

**Supp. Figure 3.**
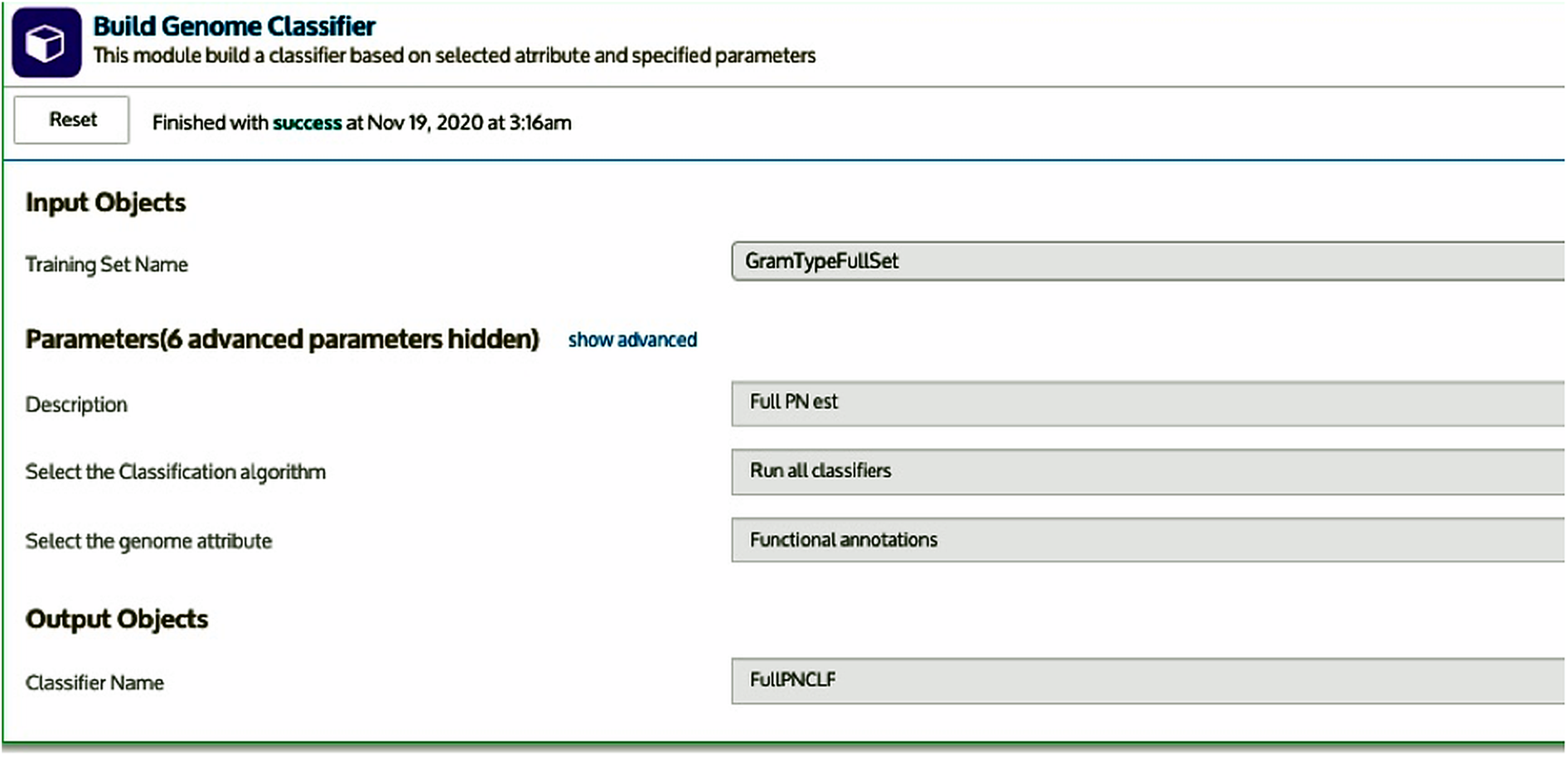
*Build Genome Classifier* app. This app takes the RAST annotated training set data that was generated from the *Annotate Genomes in a Classifier Training Set* app as an input. The app then generates six classifiers K-Nearest Neighbors (KNN), Gaussian Naive Bayes (GNB), Support Vector Machine (SVM), Neural Network (NN), Logistic Regression (LR), and Decision Tree (DT) and provides the best performing classifiers with rankings, so the user can pick the desired classifier for downstream analysis. The generated classifiers are saved in the KBase narrative as a ‘Genome Categorizer’ object. The output report generated from this app shows the confusion matrix data for each classifier algorithm, the decision tree diagram, and the highest-ranking attributes based on the decision tree.
(iv) Predict the phenotype of unclassified genomes:

**Supp. Figure 4.**
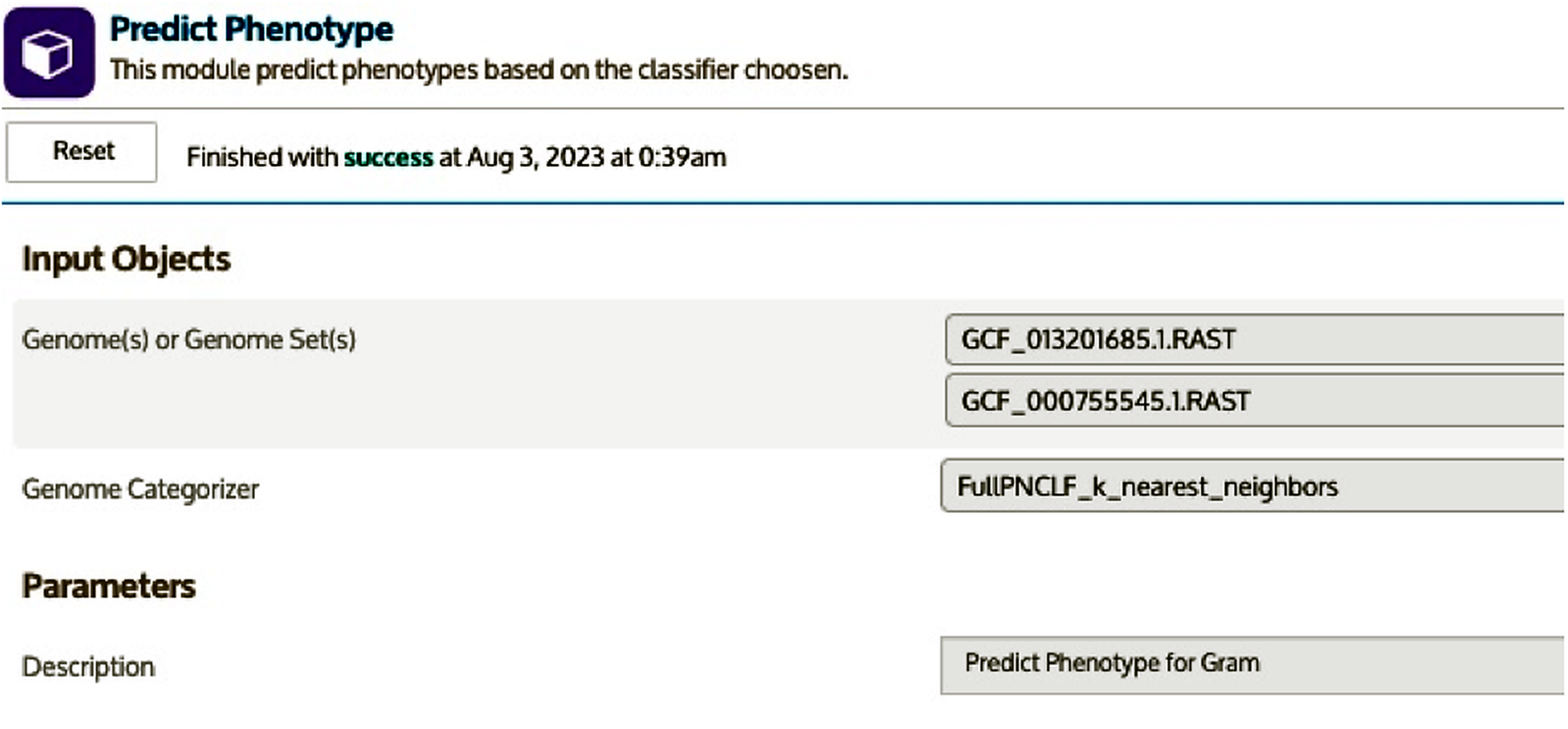
*Predict Phenotype* app. This app takes any genome or genomes as an input then classifies the organism based on the selected classifier. This app generates an output report table specifying the genomes and the predicted phenotypes.

## Acknowledgements

This work is supported as part of the BER Genomic Science Program. The DOE Systems Biology Knowledgebase (KBase) is funded by the U.S. Department of Energy, Office of Science, Office of Biological and Environmental Research under Award Numbers DE-AC02-05CH11231, DE-AC02-06CH11357, DE-AC05-00OR22725, and DE-AC02-98CH10886.

The submitted manuscript has been created by UChicago Argonne, LLC, Operator of Argonne National Laboratory (“Argonne”). Argonne, a U.S. Department of Energy Office of Science laboratory, is operated under Contract No. DE-AC02-06CH11357. The U.S. Government retains for itself, and others acting on its behalf, a paid-up nonexclusive, irrevocable worldwide license in said article to reproduce, prepare derivative works, distribute copies to the public, and perform publicly and display publicly, by or on behalf of the Government. The Department of Energy will provide public access to these results of federally sponsored research in accordance with the DOE Public Access Plan. http://energy.gov/downloads/doe-public-access-plan

